# Impaired online and enhanced offline motor sequence learning in individuals with Parkinson’s disease

**DOI:** 10.1101/2024.11.14.623568

**Authors:** Anke Van Roy, Emily Dan, Letizia Micca, Moran Gilat, Piu Chan, Julien Doyon, Genevieve Albouy, Bradley R. King

## Abstract

Whereas memory consolidation research has traditionally focused on longer temporal windows (i.e., hours to days) following an initial learning episode, recent research has also examined the functional significance of the shorter rest epochs commonly interspersed with blocks of task practice (i.e., “micro-offline” intervals on the timescale of seconds to minutes). In the motor sequence learning domain, evidence from young, healthy individuals suggests that micro-offline epochs afford a rapid consolidation process that is supported by the hippocampus. Consistent with these findings, amnesic patients with hippocampal damage were recently found to exhibit degraded micro-offline performance improvements. Interestingly, these offline losses were compensated for by larger performance gains during online practice. Given the known role of the striatum in online motor sequence learning, we hypothesized that individuals with dysfunction of the striatal system would exhibit impaired online, yet enhanced micro-offline, learning (i.e., a pattern of results opposite to those observed in patients with hippocampal lesions). We tested this hypothesis using Parkinson’s disease (PD) as a model of striatal dysfunction. Forty-two drug-naïve individuals (men and women) with a clinical diagnosis of unilateral PD and 30 healthy control subjects completed a motor sequence learning paradigm. Individuals with PD exhibited deficits during online task practice that were paralleled by greater improvements over micro-offline intervals. This pattern of results could not be explained by disease-related deficits in movement execution. These data suggest that striatal dysfunction disrupts online learning, yet total learning remains unchanged because of greater micro-offline performance improvements that potentially reflect hippocampal-mediated compensatory processes.

**Significance Statement:** The short rest intervals commonly interspersed between periods of active task engagement have traditionally been employed to minimize the build-up of fatigue. There is recent evidence, however, suggesting that these rest epochs may play an active role in motor learning and memory processes and the hippocampus appears to be a critical brain region supporting this rapid “offline” learning. Here, we show that individuals with Parkinson’s disease, a movement disorder characterized by dysfunction in the basal ganglia including the striatum, exhibit deficits during active task practice but greater learning over the interspersed offline intervals. Results potentially suggest that the relatively intact hippocampus may help compensate for motor sequence learning deficits linked to a disrupted striatal system in Parkinson’s disease.

## Introduction

Intervals of rest that follow periods of active learning are known to offer a privileged window for new information to be consolidated into stable, long-term memories [e.g., (McGaugh, 2000)]. Research examining memory consolidation processes has traditionally focused on rest intervals that span hours, days or months following initial learning (i.e., “macro-offline” intervals). However, recent research in the motor memory domain suggests that the shorter rest intervals interspersed with blocks of task practice – largely intended to minimize the accrual of fatigue - afford a “micro-offline” consolidation process that is marked by additional performance improvements (Du et al., 2016, 2017; Bönstrup et al., 2019, 2020; Jacobacci et al., 2020; Buch et al., 2021; Prashad et al., 2021; Gann et al., 2023; Mylonas et al., 2024; Van Roy et al., 2024).

At the neural level, multiple brain imaging studies have converged to indicate that micro-offline motor sequence learning (MSL) is supported by the hippocampus (Jacobacci et al., 2020; Buch et al., 2021; Gann et al., 2023; Chen et al., 2024; Sjøgård et al., 2024), and the reactivation of MSL-related patterns of hippocampal activity in particular (Buch et al., 2021; Gann et al., 2023). Consistent with these data, it was recently shown that amnesic patients with hippocampal damage present impaired micro-offline performance improvements relative to healthy, age-similar adults (Mylonas et al., 2024). Moreover, these smaller micro-offline performance improvements in amnesic patients were linked to larger online performance gains in the subsequent block of active task practice, suggesting a trade-off between online and offline performance gains (Mylonas et al., 2024).

Given that that the striatum is particularly involved in online MSL [e.g., (Lehericy et al., 2005; Albouy et al., 2013b; Andersen et al., 2020) and the striatal and hippocampal systems are known to interact during learning and memory tasks (Poldrack and Packard, 2003; Freedberg et al., 2020), including MSL (Albouy et al., 2013c, 2013b; King et al., 2017a), we hypothesized that individuals with an impairment to the striatal system would exhibit a behavioral pattern opposite to that observed in patients with an impaired hippocampus. Specifically, an affected striatal system would result in smaller online, but larger micro-offline, performance improvements. We tested this hypothesis using early Parkinson’s disease (PD) as a model of striatal dysfunction.

## Materials and Methods

This research consists of a re-analysis of the data published in (Dan et al., 2015). Complete details on the full experiment can be found in the original publication. The sections that follow provide the methodological details pertinent for the aims of the present research. Data used in the statistical analyses and depicted in the figures are included without restriction as supplemental material.

### Participants

The Research Ethics Committee of Xuanwu Hospital (Beijing, China) approved this study, and all participants provided written informed consent prior to participation. Ages of the participants ranged between 42 and 77 years, with an average age of 59.7 (SD=8.0). All participants were right-hand dominant (Oldfield, 1971), of Chinese Han ethnicity, and reported no known history of drug/alcohol abuse, psychiatric or sleep disorders. Included participants scored greater than 23 on the Mini-Mental State Examination [MMSE; (Cockrell and Folstein, 1988)]. The study sample included 44 drug-naïve individuals with a clinical diagnosis of unilateral PD as well as 30 age-similar control subjects with no known history of neurological disorders or motor impairments. The Unified Parkinson’s Disease Rating Scale and Hoehn and Yahr staging were completed by a movement disorder specialist for individuals with PD. Nineteen and 25 of the individuals with PD had motor impairments limited to the left and right hands/arms, respectively. As the motor sequence learning (MSL) task was completed with the non-dominant left hand (see below for task details), the hand used to perform the task was either affected (PD-A) or unaffected (PD-U) in these groups of individuals with PD. Although overt clinical symptoms were limited to a single upper limb, there is certainly a possibility that disease-related alterations in striatal circuitry were present – albeit to different degrees - in both striatal hemispheres at the time of testing. This would be consistent with previous research demonstrating PD-related pathology before the onset of clinical symptoms (Marsden, 1990; Fearnley and Lees, 1991; Braak et al., 2003, 2006; Siderowf and Lang, 2012) as well as earlier imaging work showing changes in cortico-striatal regions contralateral to the non-affected hand (Marek et al., 1996; Schwarz et al., 2000; Tessa et al., 2013).

Data from 2 participants (1 PD-A and PD-U) were excluded from analyses due to an inability to perform the motor sequence task (i.e., accuracy in the first session was less than 50% as well as greater than 3SD below the group mean). Table 1 contains details on the 72 participants included in analyses presented in this manuscript.

**Table 1.** Participant characteristics for the three experimental groups. PD-U = individuals with Parkinson’s disease (PD) with an unaffected tested (left) hand; PD-A = individuals with PD with an affected tested hand; Control = age-similar, healthy controls. MMSE=mini-mental state examination (Cockrell and Folstein, 1988); GDS=Geriatric depression scale (Yesavage et al., 1982); UPDRS=Unified Parkinson’s disease rating scale; L and R = left- and right-side scores of the UPDRS. Far right column provides results from statistical contrasts assessing group differences. Test for the row n / females assesses distribution of gender. * significant group differences at p < 0.05. MMSE scores were significantly higher in control as compared to both PD-U (t(52)=2.58, p=0.013) and PD-A (t(46)=2.67, p=0.010). GDS scores were lower in healthy, control participants compared to both PD-U (t(52)=2.50, p=0.016) and PD-A (t(46)=2.90, p=0.006). UPDRS III score was missing from one PD-U participant and thus this contrast one fewer degrees of freedom. As participants in the PD group exhibited unilateral symptoms, the two groups differed in the left- and right-side scores of the UPDRS.

|  | Control | PD-U | PD-A | Group Effects |
| --- | --- | --- | --- | --- |
| n / females | 30 / 16 | 24 / 12 | 18 / 9 | $\chi^2=0.08$, $p=0.96$ |
| Age (years) | 60.9±8.0 | 57.4±8.8 | 60.0±7.5 | $F_{(2,69)}=1.24$, $p=0.30$ |
| Education (years) | 11.0±2.9 | 11.0±2.9 | 9.4±3.5 | $F_{(2,69)}=1.78$, $p=0.18$ |
| MMSE | 29.1±1.1 | 28.3±1.2 | 27.9±1.9 | $F_{(2,69)}=4.84$, $p=0.01^*$ |
| GDS | 3.8±3.1 | 5.8±2.4 | 6.2±3.0 | $F_{(2,69)}=5.24$, $p=0.008^*$ |
| Disease duration (years) | - | 1.5±1.0 | 1.3±1.0 | $t_{(40)}=0.73$, $p=0.47$ |
| UPDRS I | - | 1.2±1.9 | 0.7±1.0 | $t_{(40)}=1.03$, $p=0.31$ |
| UPDRS II | - | 6.4±2.6 | 5.7±3.0 | $t_{(40)}=0.84$, $p=0.41$ |
| UPDRS III | - | 9.4±3.3 | 9.6±3.6 | $t_{(39)}=0.20$, $p=0.84$ |
| L-UPDRS III | - | 0.0±0.2 | 6.7±3.0 | $t_{(40)}=11.19$, $p<0.001^*$ |
| R-UPDRS III | - | 6.6±2.0 | 0.0±0.0 | $t_{(40)}=14.04$, $p<0.001^*$ |
| Hoehn & Yahr | - | 5.8±2.4 | 5.8±2.4 | $t_{(40)}=0.58$, $p=0.57$ |

### Motor Sequence Learning Task

The full experimental protocol consisted of two sessions separated by approximately 24 hours. In the initial visit, participants completed a series of finger tapping tasks following the administration of the clinical assessments, MMSE and GDS. For the finger tapping tasks, participants were seated in a height-adjustable chair in front of a computer monitor placed on a desk. The left (i.e., non-dominant) hand of the participants was placed on top of a four-button response box positioned just left to the midline of the participants. As a warm-up exercise, participants simultaneously pressed all four buttons with the four fingers of the left hand as fast as possible. The start of this warm-up was cued by a green fixation cross displayed on the computer monitor. Unbeknownst to the participants, the color of the cross changed to red (instructing participants to stop) following the completion of 60 presses. Participants then completed an adapted version of the motor sequence learning (MSL) task employed extensively in our team’s previous research with older adults [e.g., (Fogel et al., 2014; Dan et al., 2015; King et al., 2017b, 2020; Rumpf et al., 2017)]. In brief, participants used the four fingers of their left hand to press the corresponding buttons on the response box to perform an explicitly known sequence of finger presses: 4-1-3-2-4, where the numbers 1 through 4 represent the index, middle, ring and little fingers, respectively. Participants were first asked to perform the sequence of finger movements slowly and without any errors to verify that they memorized and could correctly perform the sequence. This phase terminated when 3 consecutive sequences were performed without an error. Subsequently, participants completed the learning phase by repeatedly performing the sequence as fast and accurately as possible while a green cross was displayed on the monitor. Participants were instructed to return to the beginning of the sequence if an error was made. The MSL included 14 practice blocks, with each block consisting of 60 key presses (ideally corresponding to 12 correct sequences). In between blocks, a red cross was displayed on the monitor for a duration of 25 seconds during which participants were instructed to rest their hand. Consistent with previous research (Fogel et al., 2014; King et al., 2017b, 2020; Rumpf et al., 2017, 2020; Van Roy et al., 2024), this task design fixed the number of movements within each practice block. This ensures the amount of motor sequence practice – quantified by the number of key presses - is the same across participants and thus experimental groups. The alternative design (i.e., fixing the duration of each block) runs the risk of affording more motor sequence practice in the groups that perform the task faster.

Participants were invited for a second experimental session that took place approximately 24 hours after the first session. During this second session, participants completed the same series of motor tasks as in the first experimental session (i.e., warm-up, verification, motor sequence training). As our primary objective was to assess micro-online and -offline learning during initial training (i.e., when performance improvements are larger), the data related to the second session are not included in the main text are contained in the Extended Data for completeness.

Performance on the motor sequence learning task was quantified in terms of speed with the outcome measure Sequence Duration (Seq Dur), defined as the time to complete a correct, 5-element sequence. [A detailed characterization of performance accuracy was included in (Dan et al., 2015)]. The mean Sequence Duration of each block was computed for each participant and used to assess performance improvements as a function of practice. Micro-*online* gains in performance were computed as the differences in Sequence Duration between the first and last correct sequences within each block (i.e., first correct sequence of block *n* – last correct sequence of block *n*). Micro-*offline* performance gains were computed as the differences in Sequence Duration between the last correct sequence of one block and the first correct sequence of the subsequent block (i.e., last correct sequence block *n* – first correct sequence block *n+1*). Micro-online performance gains for a particular block plus the micro-offline gain over the subsequent rest interval were summed to compute a total gain score. Micro-online, micro-offline and total performance gains were summed across blocks and subjected to the statistical analyses described below. For all three metrics, higher values are indicative of greater performance improvements in speed.

### Statistical Analyses

Statistical analyses were conducted in RStudio version 2023.09.01 (Posit, PBC, Boston, Massachusetts, USA). All null hypothesis statistical tests were two-sided and considered significant when p<0.05. If the sphericity assumption was violated, a Greenhouse-Geisser correction was applied. For any main effects of Group, pairwise (i.e., between pairs of experimental groups) comparisons were conducted with corrections for multiple comparisons using the false discovery rate (FDR) method (Benjamini and Hochberg, 1995). Depending on the statistical tests, generalized eta-squared (ges) or Hedges’ g were reported as effect size measures. The correspondence among reported p-values, test statistics and degrees of freedom was verified via statcheck (Epskamp and Nuijten, 2018). Null hypothesis statistical testing was complemented with the computation of Bayes Factors (BF) using the anovabf and ttestbf functions in R. BFs reflect the likelihood that the observed data favor an alternative model (e.g., evidence for differences among groups) relative to the null (e.g., evidence for no differences). For mixed ANOVAs, the null models were specified as the random effect of subjects. For one-way ANOVAs, the null models included only the intercept term. For t-tests, the null model was specified as the difference between means equal to zero. Default parameters in the R functions were used, with the exception that “whichModels” was set to “all” for mixed model ANOVAs to extract the appropriate BFs. Our results section reports BF_10_ values, with larger values indicative of greater likelihood that the observed data favor the alternative as compared to the null hypotheses.

Group differences in Sequence Duration were assessed using a mixed ANOVA, with Block (14 levels) as the within-subjects and Group (3 levels: Control, PD-A and PD-U) as the between-subjects factor. Group differences in total performance gains were assessed with a one-way ANOVA with Group (3 levels) as the between-subjects factor. Performance changes over micro intervals were first assessed with a Group (3 levels) by Interval (online vs. offline) ANOVA. A significant Group x Interval interaction was followed up with separate one-way ANOVAs (between-subject factor Group) on micro-online and -offline performance gains.

## Results

The average time to complete a correct sequence (Sequence Duration (or Seq Dur); Figure 1A) significantly decreased as a function of practice (effect of block: F_(3.76,259.19)_=58.75, p<0.001, ges=0.119, BF_10_=8.24e^111^). These changes in performance across blocks did not statistically differ among the three groups (group by block interaction: F_(7.51,259.19)_=0.66, p=0.71, ges=0.003, BF_10_=0.006). As expected, the PD-A group was slower, on average, to complete correct motor sequences as compared to both the controls and the PD-U group (Main effect of Group: F_(2,69)_=5.28, p=0.007, ges=0.114, BF_10_=6.40; PD-A vs. controls: F_(1,46)_=11.62, p=0.001, FDR-corrected p=0.003, ges=0.169, BF_10_=23.64; PD-A vs. PD-U: F_(1,40)_=4.97, p=0.031, FDR-corrected p=0.047, ges=0.097, BF_10_=2.17; PD-U vs. controls: F_(1,52)_=0.58, p=0.45, ges=0.009, BF_10_=0.50).

**Figure 1.**
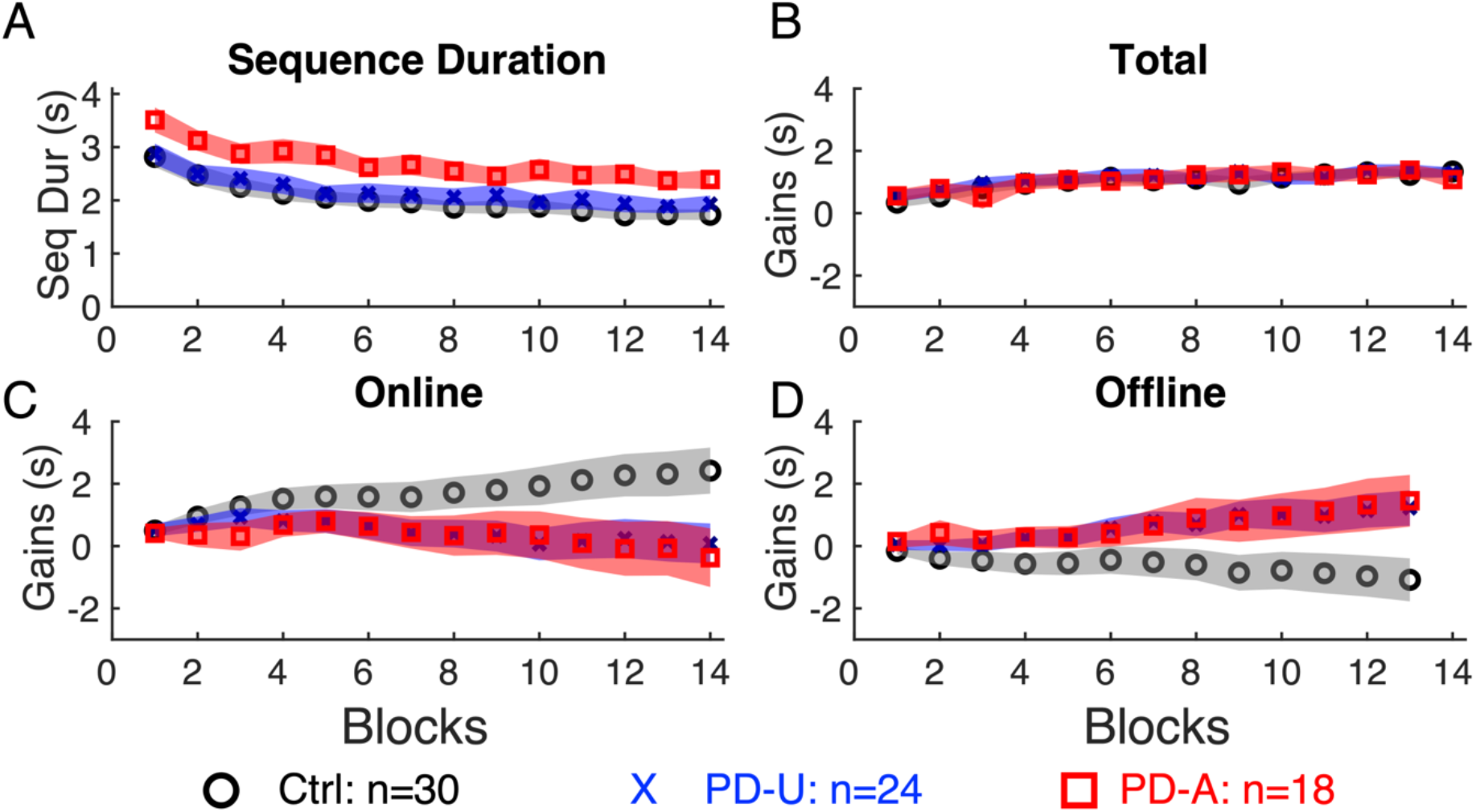
**A**. Mean sequence Duration (Seq Dur), defined as the time to complete a correct 5-element sequence of finger movements with the non-dominant left hand, is plotted as a function of practice blocks during an initial training session for the control participants (black circles) and individuals with Parkinson’s disease with an affected (PD-A; red squares) or unaffected tested (left) hand (PD-U; blue crosses). Although the PD-A group was slower on average, the three groups learned the motor sequence and to a similar extent (see main text for statistical information). **B-D**. Cumulative sums of total (Panel B), micro-online (Online; Panel C) and micro-offline (Offline; Panel D) performance gains for the three groups. For all 3 metrics, positive values are indicative of performance improvements. As the training session consisted of 14 practice blocks, there were 13 inter-block intervals for the assessment of Offline performance gains. The final cumulative sums of total, Online and Offline gains are depicted in Figure 2 with the corresponding statistical analyses presented in the main text. For all 4 panels, data points represent group- and block-specific averages, and the shaded regions depict standard error of the mean. See Extended Data Figure 1-1 for a depiction of Sequence Duration of the first and last correct sequences – used in the computation of micro-online and -offline performance changes, within each practice block.

These data demonstrate that the PD-A group, but not the PD-U group, exhibited deficits in the *execution* of motor sequences with the left hand. However, there was no credible evidence to suggest that the *learning* of the motor sequence, as assessed by changes in performance across blocks of practice, was impacted in either PD group.

We then assessed the effects of PD on micro-online (i.e., performance changes within blocks of practice), micro-offline (i.e., performance changes between consecutive practice blocks) as well as total performance gains (i.e., aggregate of micro-online and -offline gains). Panels B-D in Figure 1 depict cumulative sums of these three performance gains as a function of practice blocks (i.e., values at block *n* show the across-participant average of the sums of the gain values from block 1 through block *n*). Visual inspection of these data suggests similar total learning gains, but differences among groups in the relative contributions of micro-online and -offline performance improvements.

Statistical analyses were conducted on the final cumulative sums (Figure 2). Total learning gains were significantly greater than zero in each of the three groups (see Table 2-1 in Extended Data for details from within-group statistical contrasts) and the magnitudes of the total performance gains did not statistically differ among the three groups (F_(2,69)_=0.45, p=0.64, ges=0.013, BF_10_=0.17; Figure 2A). These data again suggest similar total learning among the 3 groups.

**Figure 2.**
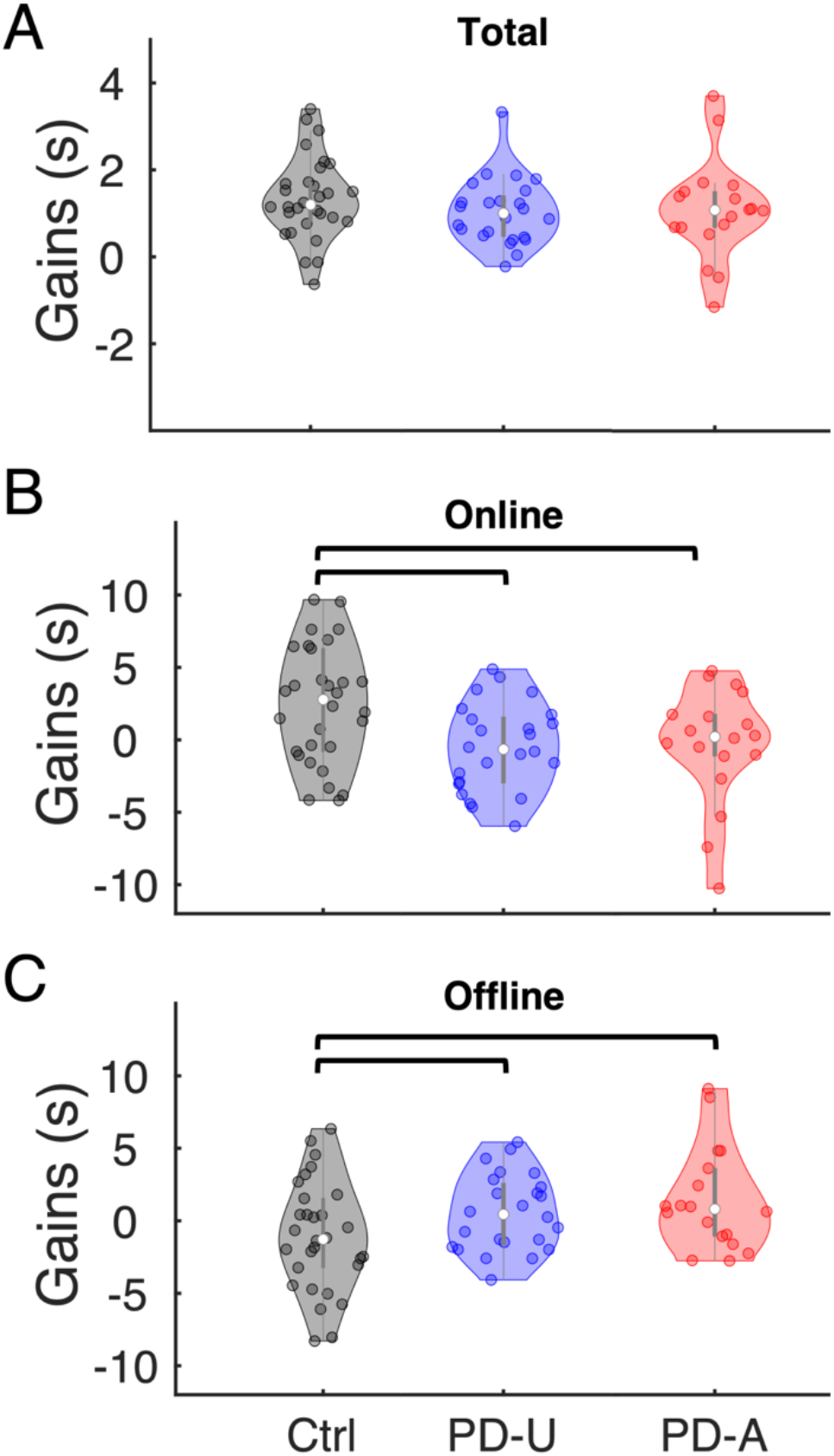
Total (Panel A), micro-online (Online, B) and micro-offline (Offline, B) performance gains summed across blocks during an initial training session for the control (black), PD-U (blue) and PD-A (red) groups. Positive values of all 3 metrics are indicative of performance improvements. Shaded regions represent kernel density estimates of the data, colored circles depict individual data, open circles represent group medians and vertical gray lines show interquartile ranges (Bechtold, 2016). Horizontal lines with brackets: p < 0.05. The primary findings remained the same when age, gender and education levels were separately included as a covariate in the statistical model (see Extended Data Table 2-2). Extended Data Table 2-1 contains statistical output from the within-group statistical contrasts of the depicted total, online and offline gains. Figure 2-1 and Table 2-3 in Extended Data contain analogous results pertaining to performance in the 24-hour retest session. See control analyses presented in the main text and Figure 2-2 in Extended Data for results with an alternative computation for micro-offline performance gains that minimizes the effect of reactive inhibition that may accrue during blocks of active task practice.

Micro-online and -offline gains were first subjected to a Group (3 levels) x Interval (2 levels: online vs. offline) ANOVA and results revealed a significant interaction (F_(2,69)_=4.25, p=0.018, ges=0.108, BF_10_=146.7). Follow-up simple effect tests revealed that the three groups differed in the magnitude of both micro-online and -offline performance gains (micro-online: F_(2,69)_=4.03, p=0.022, ges=0.105, BF_10_=2.60, Figure 2B; micro-offline: F_(2,69)_=4.37, p=0.016, ges=0.113, BF_10_=3.38, Figure C). Specifically, and as hypothesized, both the PD-U and the PD-A groups exhibited significantly *smaller* micro-online gains as compared to the controls (PD-U vs. controls: t_(52)_=2.31, p=0.025, FDR-corrected p=0.037, Hedges g=0.62, BF_10_=2.39; PD-A vs. controls: t_(46)_=2.32, p=0.025, FDR-corrected p=0.037, Hedges g=0.68, BF_10_=2.46; PD-U vs. PD-A: t_(40)_=0.41, p=0.68, Hedges g=0.13, BF_10_=0.33). Conversely, both the PD-U and PD-A groups exhibited significantly *larger* micro-offline gains as compared to controls (PD-U vs. controls: t_(52)_=2.50, p=0.016, FDR-corrected p=0.038, Hedges g=0.67, BF_10_=3.37; PD-A vs. controls: t_(46)_=2.31, p=0.026, FDR-corrected p=0.038, Hedges g=0.68, BF_10_=2.39; PD-U vs. PD-A: t_(40)_=0.24, p=0.81, Hedges g=0.07, BF_10_=0.31). These results demonstrate that individuals with PD exhibited impaired micro-online and enhanced micro-offline motor learning as compared to age-similar controls.

Ideally, the first and last correct sequences – used in the quantification of micro-online and -offline performance changes - would consist of key presses 1-5 and 56-60, respectively. However, any incorrect key presses may shift the position of the first and last correct sequences within a given practice block. We extracted the mean indices of the first and last correct sequences within each block and then assessed potential group differences with a one-way ANOVA (between-subject factor Group: 3 levels). On average, the first press of the first correctly performed sequence was key 2.20. That is, the first correct sequence included key presses ∼2-6. Similarly, the last correctly performed sequence, on average, included key presses ∼53-57 (out of 60). Importantly, there were no differences among the experimental groups in the indices of the first and last correct sequences (First: F_(2,69)_=1.63, p=0.20, ges=0.045, BF_10_=0.43; Last: F_(2,69)_=2.35, p=0.10, ges=0.064, BF_10_=0.73). Furthermore, to assess whether the positions of the first and last correct sequences influenced the assessment of group differences in micro-online and -offline performance gains, separate ANCOVAs – with the indices of the first and last correct sequences as a covariate - were conducted on each gain metric. The significant group differences in the magnitude of micro-online and -offline gains reported above remained significant when the indices of the first and last correct sequences were included as a covariate in the statistical models (Group main effects: all F_(2,68)_ > 3.51; all p values < 0.036). These data collectively suggest that the between-group differences in micro-online and -offline performance changes cannot be explained by differences in the positions of the first and last correct sequences within practice blocks.

It has been argued that the micro-offline performance gains reflect – at least partially - the dissipation of reactive inhibition that accrues over the previous online epoch (Gupta and Rickard, 2022, 2024). Reactive inhibition is defined as the slowing of movement during repetitive task practice that may be the result of one or multiple factors such as fatigue, inattention, decreased motivation, etc. In the context of the current results, one could speculate that the accrual of reactive inhibition is disproportionately larger in patients with PD as compared to healthy controls. To address this possibility, and as done in previous research [e.g., (Du et al., 2017)], micro-offline gains were re-computed relative to the fastest – as opposed to the last – correctly performed sequence within each practice block. Computing gains relative to the fastest correctly performed sequence thus serves to minimize the effects of reactive inhibition that may accrue within blocks of practice. These adjusted micro-offline performance gains were then subjected to a one-way ANOVA with Group as the between-subjects factor. Results revealed a significant main effect of Group (see Figure 2-2 in Extended Data; micro-offline: F_(2,69)_=4.46, p=0.015, ges=0.115, BF_10_=3.63). Follow-up pairwise comparisons revealed that the PD-U and the PD-A groups exhibited significantly *larger* micro-offline gains as compared to the controls (PD-U vs. controls: t_(52)_=2.59, p=0.013, FDR-corrected p=0.038, Hedges g=0.70, BF_10_=4.01; PD-A vs. controls: t_(46)_=2.27, p=0.028, FDR-corrected p=0.042, Hedges g=0.66, BF_10_=2.22; PD-U vs. PD-A: t_(40)_=0.09, p=0.93, Hedges g=0.03, BF_10_=0.31). This suggests that the larger micro-offline gains observed in individuals with PD cannot be explained by a disproportionately larger worsening of performance within a block due to the accrual of reactive inhibition that then dissipates during the micro-offline epochs.

## Discussion

The current results demonstrate that individuals with PD exhibit impaired motor sequence learning during online task practice, yet these deficits are paralleled by greater performance gains during the interleaved offline epochs. As the groups of individuals with PD were limited to individuals recently diagnosed and with Hoehn and Yahr stages of 1 or 1.5, these results suggest that this pattern of disrupted online but enhanced offline learning appears relatively early in the progression of the disease.

Interestingly, the impaired online but enhanced offline learning exhibited by the individuals with PD cannot be attributed to disease-related deficits in movement execution, as the same effects were also observed in the PD-U group with no clinically observable movement deficits in the (left) hand used to complete the motor sequence task. The lack of deficits in the tested hand of the PD-U group was further evidenced by our assessment of group differences in the average time to complete movement sequences (i.e., Figure 1A). The lack of differences between the two groups of individuals with PD is also interesting in the context of disease progression. Specifically, even though overt clinical symptoms were limited to a single limb in the individuals with PD, it is likely that disease-related alterations were present in both striatal hemispheres at the time of testing. In this context, it is possible that the underlying pathology in the PD-U cohort was not sufficiently advanced to result in symptoms during clinical testing but was advanced enough to reveal altered online and offline motor sequence learning. On the one hand, this raises the intriguing possibility that the pattern of disrupted online yet enhanced offline learning may appear prior to the onset of any clinical symptom (i.e., in either limb) and thus these behavioral metrics may hold some value as a non-invasive, biomarker for early identification of PD. On the other hand, there is considerable inter-individual variability in these metrics and the large overlap among the distributions from the three experimental groups (see Figure 2) dampens enthusiasm for such markers to be used – at the individual level – for the purpose of early identification.

The results of this research on individuals with PD can be contrasted with the recent work in amnesic patients with hippocampal damage that demonstrated smaller and larger offline and online performance gains, respectively (Mylonas et al., 2024). Bridging these two studies, results suggest that: a) impairments to the striatum or the hippocampus result in performance deficits during micro-online or -offline epochs, respectively; and) these deficits are mirrored by greater performance gains over the alternate intervals. These data may then suggest that impairment to one region triggers compensatory processes in the other region, affording unimpaired total learning in both clinical populations. Specifically, these results can be interpreted in the context of previously posited frameworks that speak to the interaction between striatal and hippocampal systems during learning, and motor sequence learning in particular [e.g., (Albouy et al., 2013b; Freedberg et al., 2020)]. In brief, hippocampo-cortical and striato-cortical systems are thought to competitively interact during initial sequence learning. This interaction is potentially mediated by the prefrontal cortex and - at least in a healthy, unimpaired system - is thought to afford the development of a memory trace that includes at least two representations of the acquired motor sequence: a) a hippocampal-mediated spatial representation; and, b) a striatal-mediated motoric representation (Albouy et al., 2013b, 2015). In the context of the current results, it could be speculated that the smaller online performance gains observed in the individuals with PD reflects the disruption of the striatal-mediated, motoric sequence representation typically developed as a function of online task practice. Despite these online performance losses, individuals with PD still exhibit comparable sequence learning because of larger micro-offline improvements. We posit that this enhanced offline learning in individuals with PD can be attributed to an upweighting, perhaps orchestrated via the prefrontal cortex, of the intact hippocampal-cortical system which favored the development of the hippocampal-mediated sequence representation. Extending this framework to the aforementioned research in amnesic patients (Mylonas et al., 2024), we speculate that hippocampal damage in these individuals disrupted the hippocampal-mediated sequence representation, compromising micro-offline performance improvements. A shift in balance to the unimpaired striatal system in these patients might have favored the development of the striatal-mediated sequence representation, which was reflected by larger online performance gains. One area of future research is to empirically test these difference sequence representations – as in (Albouy et al., 2013a, 2015) – in individuals with compromised hippocampal and/or striatal systems.

It is worth noting that there is currently a discussion in the literature as to whether micro-offline performance improvements reflect an active, consolidation-like mnemonic process (Gupta and Rickard, 2022, 2024; Das et al., 2024). An alternative explanation is that micro-offline performance improvements can be explained by the dissipation of reactive inhibition (Gupta and Rickard, 2022, 2024). Specifically, reactive inhibition, defined as a slowing of movement during continuous motor task practice (i.e., potentially due to one or multiple non-learning-related factors such as fatigue, inattention, etc.), accrues over the course of an online practice block and then dissipates during the rest periods interleaved with task practice. In the context of the current research, one could argue that reactive inhibition is greater in individuals with PD and thus the larger micro-offline performance gains in the PD groups simply reflect the dissipation of this greater reactive inhibition. While we cannot completely discount the effects of reactive inhibition, this framework is not fully consistent with our data as well recent findings reported in the literature. For example, we re-computed micro-offline gains to assess changes in performance from the fastest – as compared to the last – correctly performed sequence in one block to the first correct sequence in the subsequent block. This computation minimizes – although certainly does not eliminate - the effects of reactive inhibition that may accrue over the course of a given block. Results indicated that these adjusted micro-offline performance gains were also significantly larger in individuals with PD as compared to healthy controls (see control analyses in results section and Figure 2-2 in Extended Data). Second, it is difficult to reconcile the reactive inhibition framework with the neuroimaging results in the available literature. Specifically, the replay of learning-related patterns of brain activity during micro-offline intervals (Buch et al., 2021; Gann et al., 2023) and the link between hippocampal ripples and micro-offline performance gains (Chen et al., 2024; Sjøgård et al., 2024) suggest an active consolidation process that is incompatible with the reactive inhibition explanation. Nonetheless, these conflicting explanations highlight the fact that causal, experimental evidence differentiating these interpretations is currently lacking and warrants attention in future research.

In conclusion, our results demonstrate that individuals with Parkinson’s disease exhibit impaired online, yet enhanced offline, motor sequence learning. These results cannot be attributed to deficits in movement execution and do not appear to be solely linked to the accrual of reactive inhibition during motor task practice. Rather, we suggest that striatal dysfunction as a result of PD specifically disrupts online learning processes, yet total learning remains relatively unchanged due to greater offline improvements during the interspersed rest periods. We speculate that this enhanced micro-offline performance improvements in individuals with PD is supported by compensatory processes supported by the relatively intact hippocampus. It should be explicitly stated that our sample size cannot be considered large and thus similar analyses should be conducted on larger participant samples in future research. Similarly, it would be interesting to assess micro-online and -offline performance gains in participants with more advanced disease stages than tested in the current research. One could speculate that the disrupted online and enhanced offline learning evident in PD would increase as a function of disease progression. This hypothesis awaits investigation.

## Conflicts of Interest

None

## Acknowledgements

The original study (Dan et al., 2015; *PLoS One*) was supported by grants from the Ministry of Science and Technology of China (2012AA02A514), the National Basic Research Development Program of China (2011CB504101), Ministry of Health (201002011), Beijing High Standard Health Human Resource Cultural Program in Health System (2009e1e12) awarded to P.C. as well as the European Commission – ERA Net Neuron (Identifying #26659) awarded to J.D.

## Notes

### Competing Interest Statement

The authors have declared no competing interest.

### Summary of Updates

1) While revisiting the participant characteristic information, we noticed that there was one healthy control participant that was (by mistake) not included in the original version of this manuscript. We apologize for this error. We have now included this participant in all analyses. Accordingly, statistical information (i.e., test statistics, p values, degrees of freedom, effect sizes and Bayes Factors) has been updated in the revised version. Importantly, the findings of this research remain unchanged. 2) The revised version has an expanded discussion on the lack of differences between the two PD cohorts as well as the possible compensatory interaction between striatal and hippocampal systems. 3) Table 1 with participant characteristic information has been added. 4) Minor edits were made in the revised version.

## References

Albouy G, Fogel S, King BR, Laventure S, Benali H, Karni A, Carrier J, Robertson E, Doyon J (2015) Maintaining vs. enhancing motor sequence memories: Respective roles of striatal and hippocampal systems. NeuroImage 108:423–434.

Albouy G, Fogel S, Pottiez H, Nguyen VA, Ray L, Lungu O, Carrier J, Robertson E, Doyon J (2013a) Daytime sleep enhances consolidation of the spatial but not motoric representation of motor sequence memory. PloS One 8:e52805–e52805.

Albouy G, King BR, Maquet P, Doyon J (2013b) Hippocampus and striatum: Dynamics and interaction during acquisition and sleep-related motor sequence memory consolidation. Hippocampus 23:985–1004.

Albouy G, Sterpenich V, Vandewalle G, Darsaud A, Gais S, Rauchs G, Desseilles M, Boly M, Dang-Vu T, Balteau E, Degueldre C, Phillips C, Luxen A, Maquet P (2013c) Interaction between hippocampal and striatal systems predicts subsequent consolidation of motor sequence memory. PloS One 8:e59490–e59490.

Andersen KW, Madsen KH, Siebner HR (2020) Discrete finger sequences are widely represented in human striatum. Scientific Reports 10:1–12.

Bechtold B (2016) Violin Plots for Matlab. Github Project https://github.com/bastibe/Violinplot-Matlab, DOI: 10.5281/zenodo4559847.

Benjamini Y, Hochberg Y (1995) Controlling the False Discovery Rate: A Practical and Powerful Approach to Multiple Testing. Journal of the Royal Statistical Society: Series B (Methodological) 57:289–300.

Bönstrup M, Iturrate I, Hebart MN, Censor N, Cohen LG (2020) Mechanisms of offline motor learning at a microscale of seconds in large-scale crowdsourced data. Science of Learning 5:7–7.

Bönstrup M, Iturrate I, Thompson R, Cruciani G, Censor N, Cohen LG (2019) A Rapid Form of Offline Consolidation in Skill Learning. Current Biology 29:1346-1351.e4.

Braak H, Bohl JR, Müller CM, Rüb U, de Vos RAI, Del Tredici K (2006) Stanley Fahn Lecture 2005: The staging procedure for the inclusion body pathology associated with sporadic Parkinson’s disease reconsidered. Movement Disorders 21:2042–2051.

Braak H, Del Tredici K, Rüb U, de Vos RAI, Jansen Steur ENH, Braak E (2003) Staging of brain pathology related to sporadic Parkinson’s disease. Neurobiology of Aging 24:197– 211.

Buch E, Claudino L, Quentin R, Bönstrup M, Cohen LG (2021) Consolidation of human skill linked to waking hippocampo-neocortical replay. Cell Reports.

Chen P-C, Stritzelberger J, Walther K, Hamer H, Staresina B (2024) Hippocampal ripples during offline periods predict human motor sequence learning. :2024.10.06.614680 Available at: https://www.biorxiv.org/content/10.1101/2024.10.06.614680v1 [Accessed October 9, 2024].

Cockrell JR, Folstein MF (1988) Mini-Mental State Examination (MMSE). Psychopharmacology Bulletin 24:689–692.

Dan E, King BR, Doyon J, Chan P (2015) Motor sequence learning and consolidation in unilateral de novo patients with Parkinson’s disease. PloS One 10:e0134291–e0134291.

Das A, Karagiorgis A, Diedrichsen J, Stenner M-P, Azanon E (2024) “Micro-offline gains” convey no benefit for motor skill learning. :2024.07.11.602795 Available at: https://www.biorxiv.org/content/10.1101/2024.07.11.602795v2 [Accessed October 28, 2024].

Du Y, Prashad S, Schoenbrun I, Clark JE (2016) Probabilistic Motor Sequence Yields Greater Offline and Less Online Learning than Fixed Sequence. Front Hum Neurosci 10 Available at: https://www.frontiersin.org/journals/human-neuroscience/articles/10.3389/fnhum.2016.00087/full [Accessed October 9, 2024].

Du Y, Valentini NC, Kim MJ, Whitall J, Clark JE (2017) Children and Adults Both Learn Motor Sequences Quickly, But Do So Differently. Frontiers in Psychology 08:158–158.

Epskamp S, Nuijten MB (2018) Statcheck: extract statistics from articles and recompute p-values. Available at: https://statcheck.io.

Fearnley JM, Lees AJ (1991) Ageing and Parkinson’s disease: substantia nigra regional selectivity. Brain 114:2283–2301.

Fogel SM, Albouy G, Vien C, Popovicci R, King BR, Hoge R, Jbabdi S, Benali H, Karni A, Maquet P, Carrier J, Doyon J (2014) fMRI and sleep correlates of the age-related impairment in motor memory consolidation. Human Brain Mapping 35:3625–3645.

Freedberg M, Toader AC, Wassermann EM, Voss JL (2020) Competitive and cooperative interactions between medial temporal and striatal learning systems. Neuropsychologia 136:107257–107257.

Gann MA, Dolfen N, King BR, Robertson EM, Albouy G (2023) Prefrontal stimulation as a tool to disrupt hippocampal and striatal reactivations underlying fast motor memory consolidation. Brain Stimulation 16:1336–1345.

Gupta MW, Rickard TC (2022) Dissipation of reactive inhibition is sufficient to explain post-rest improvements in motor sequence learning. NPJ Sci Learn 7:25.

Gupta MW, Rickard TC (2024) Comparison of online, offline, and hybrid hypotheses of motor sequence learning using a quantitative model that incorporate reactive inhibition. Sci Rep 14:4661.

Jacobacci F, Armony JL, Yeffal A, Lerner G, Amaro E, Jovicich J, Doyone J, Della-Maggiore V (2020) Rapid hippocampal plasticity supports motor sequence learning. Proceedings of the National Academy of Sciences of the United States of America 117:23898–23903.

King BR, Hoedlmoser K, Hirschauer F, Dolfen N, Albouy G (2017a) Sleeping on the motor engram: The multifaceted nature of sleep-related motor memory consolidation. Neuroscience & Biobehavioral Reviews 80:1–22.

King BR, Rumpf JJ, Heise KF, Veldman MP, Peeters R, Doyon J, Classen J, Albouy G, Swinnen SP (2020) Lateralized effects of post-learning transcranial direct current stimulation on motor memory consolidation in older adults: An fMRI investigation. NeuroImage 223:117323–117323.

King BR, Saucier P, Albouy G, Fogel S, Rumpf J, Klann J, Buccino G, Binkofski F, Classen J, Karni A, Doyon J (2017b) Cerebral activation during initial motor learning forecasts subsequent sleep-facilitated memory consolidation in older adults. Cerebral Cortex 27:1588–1601.

Kish SJ, Shannak K, Hornykiewicz O (1988) Uneven pattern of dopamine loss in the striatum of patients with idiopathic Parkinson’s disease. Pathophysiologic and clinical implications. The New England Journal of Medicine 318:876–880.

Lehericy S, Benali H, Van de Moortele PF, Pelegrini-Issac M, Waechter T, Ugurbil K, Doyon J (2005) Distinct basal ganglia territories are engaged in early and advanced motor sequence learning. Proceedings of the National Academy of Sciences of the United States of America 102:12566–12571.

Marek KL, Seibyl JP, Zoghbi SS, Zea-Ponce Y, Baldwin RM, Fussell B, Charney DS, van Dyck C, Hoffer PB, Innis RP (1996) [123I] beta-CIT/SPECT imaging demonstrates bilateral loss of dopamine transporters in hemi-Parkinson’s disease. Neurology 46:231–237.

Marsden CD (1990) Parkinson’s disease. Lancet 335:948–952.

McGaugh JL (2000) Memory--a century of consolidation. Science 287:248–251.

Mylonas D, Schapiro AC, Verfaellie M, Baxter B, Vangel M, Stickgold R, Manoach DS (2024) Maintenance of Procedural Motor Memory across Brief Rest Periods Requires the Hippocampus. J Neurosci 44 Available at: https://www.jneurosci.org/content/44/14/e1839232024 [Accessed September 12, 2024].

Oldfield RC (1971) The assessment and analysis of handedness: the Edinburgh inventory. Neuropsychologia 9:97–113.

Poldrack RA, Packard MG (2003) Competition among multiple memory systems: converging evidence from animal and human brain studies. Neuropsychologia 41:245–251.

Prashad S, Du Y, Clark JE (2021) Sequence structure has a differential effect on underlying motor learning processes. Journal of Motor Learning and Development 9:38–57.

Rumpf JJ, May L, Fricke C, Classen J, Hartwigsen G (2020) Interleaving Motor Sequence Training with High-Frequency Repetitive Transcranial Magnetic Stimulation Facilitates Consolidation. Cerebral Cortex 30:1030–1039.

Rumpf J-J, Wegscheider M, Hinselmann K, Fricke C, King BR, Weise D, Klann J, Binkofski F, Buccino G, Karni A, Doyon J, Classen J (2017) Enhancement of motor consolidation by post-training transcranial direct current stimulation in older people. Neurobiology of Aging 49:1–8.

Schwarz J, Linke R, Kerner M, Mozley PD, Trenkwalder C, Gasser T, Tatsch K (2000) Striatal dopamine transporter binding assessed by [I-123]IPT and single photon emission computed tomography in patients with early Parkinson’s disease: implications for a preclinical diagnosis. Archives of Neurology 57:205–208.

Siderowf A, Lang AE (2012) Premotor Parkinson’s disease: concepts and definitions. Movement Disorders 27:608–616.

Sjøgård M, Baxter B, Mylonas D, Thompson M, Kwok K, Driscoll B, Tolosa A, Shi W, Stickgold R, Vangel M, Chu CJ, Manoach DS (2024) Hippocampal ripples predict motor learning during brief rest breaks in humans. :2024.05.02.592200 Available at: https://www.biorxiv.org/content/10.1101/2024.05.02.592200v2 [Accessed March 6, 2025].

Tessa C, Diciotti S, Lucetti C, Baldacci F, Cecchi P, Giannelli M, Bonuccelli U, Mascalchi M (2013) fMRI changes in cortical activation during task performance with the unaffected hand partially reverse after ropinirole treatment in de novo Parkinson’s disease. Parkinsonism & Related Disorders 19:265–268.

Van Roy A, Albouy G, Burns RD, King BR (2024) Children exhibit a developmental advantage in the offline processing of a learned motor sequence. Commun Psychol 2:1–13.

Yesavage JA, Brink TL, Rose TL, Lum O, Huang V, Adey M, Leirer VO (1982) Development and validation of a geriatric depression screening scale: a preliminary report. Journal of Psychiatric Research 17:37–49.

