## Extended Data for "Impaired online and enhanced offline motor sequence learning in individuals with Parkinson’s disease"

**Abbreviated Title**: Rapid Consolidation in Parkinson's Disease

**Authors**: Anke Van Roy[1], Emily Dan[2,3], Letizia Micca[4], Moran Gilat[4], Piu Chan[2,3,5,6], Julien Doyon[7], Genevieve Albouy[1,8,9], Bradley R. King[1*]

[1] Department of Health & Kinesiology, University of Utah, Salt Lake City, Utah, USA

[2] Department of Neurology, Xuanwu Hospital of Capital Medical University, Beijing, China.

[3] Key Laboratory on Neurodegenerative Disorders of Ministry of Education, Key Laboratory on Parkinson's Disease of Beijing, Beijing, China.

[4] Research Group for Neurorehabilitation (eNRGy), Department of Rehabilitation Sciences, KU Leuven, Leuven, Belgium

[5] Department of Neurobiology, Xuanwu Hospital of Capital Medical University, Beijing, China.

[6] Department of Neurosurgery, Xuanwu Hospital of Capital Medical University, Beijing, China.

[7] McConnell Brain Imaging Centre, Department of Neurology and Neurosurgery, Montreal Neurological Institute, McGill University, Montreal, Canada

[8] Department of Movement Sciences, Movement Control and Neuroplasticity Research Group, KU Leuven, Leuven, Belgium

[9] LBI - KU Leuven Brain Institute, KU Leuven, Leuven, Belgium

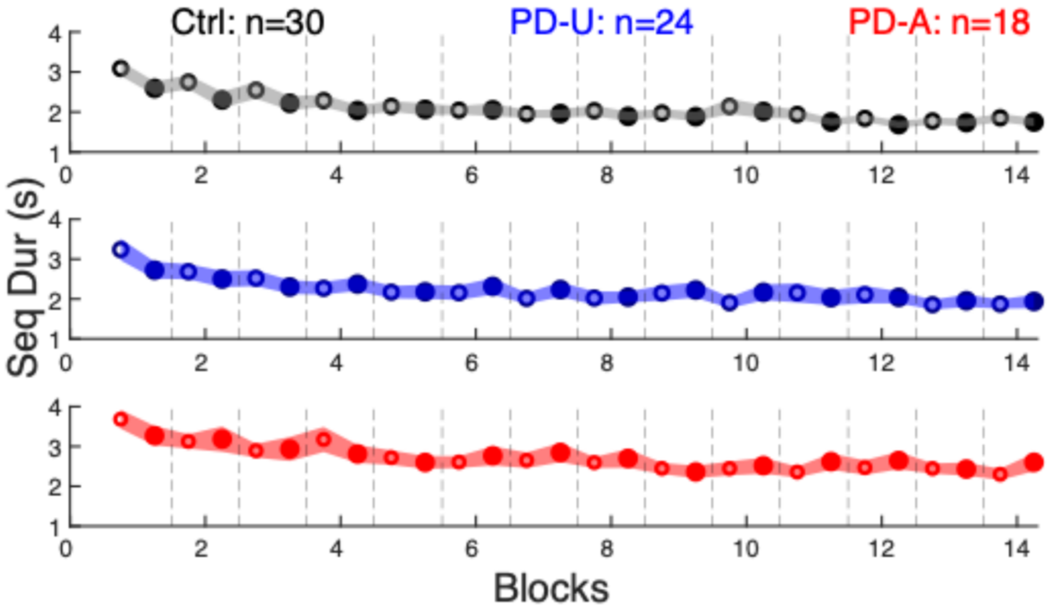


*Figure 1-1. Sequence Duration (Seq Dur) for the first (open circles) and last (closed circles) correct sequences within each practice block of an initial training session are plotted for the control participants (black; top panel) and individuals with Parkinson's disease with an affected (PD-A; red) or unaffected left hand (PD-U; blue) used to perform the task. Data points represent group- and block-specific averages, and the shaded regions depict standard error of the mean. Dashed vertical lines serve as demarcations between blocks. Sequence Durations from the first and last correct sequences within practice blocks are used to compute micro-online and -offline performance changes (see Figures 1 and 2 in the main text).*

*Table 2-1. Within-group statistical results from the initial learning session.*

| | Ctrl | PD-U | PD-A |
|---|---|---|---|
| Total | **$t_{(29)}$=7.81, p<0.001** | **$t_{(23)}$=7.98, p<0.001** | **$t_{(17)}$=4.00 p<0.001** |
| Online | **$t_{(29)}$=3.26, p=0.003** | $t_{(23)}$=0.12, p=0.91 | $t_{(17)}$=0.40, p=0.70 |
| Offline | $t_{(29)}$=1.57, p=0.13 | **$t_{(23)}$=2.16, p=0.042** | $t_{(17)}$=1.75, p=0.099 |

*Statistical output of one-sample t-tests assessing whether the magnitudes of total, micro-online (Online) and micro-offline (Offline) gains in performance from an initial training session differed from a test value of 0. Bold = significant at $p < 0.05$. All three groups exhibited positive total performance gains that significantly differed from 0, suggesting learning of the motor sequence. The magnitude of micro-online learning was significantly positive in the control participants. Conversely, micro-offline gains were significantly positive in the individuals with Parkinson's disease with an unaffected tested (left) hand (PD-U). An analogous finding was observed in individuals with Parkinson's disease with an affected tested hand (PD-A), but it is worth noting that this effect exhibited only a trend for significance (i.e., $p < 0.1$). Consistent with the between-group comparisons presented in the main text (see Figure 2), these results collectively suggest that motor sequence learning is largely achieved via online and offline performance improvements in control participants and individuals with PD, respectively.*

*Table 2-2. Group differences in performance metrics when controlling for potential confounding factors.*

| | Age | Gender | Education |
|---|---|---|---|
| Total | $F_{(2,68)}$=0.44, p=0.65 | $F_{(2,68)}$=0.43, p=0.65 | $F_{(2,68)}$=0.67, p=0.52 |
| Online | **$F_{(2,68)}$=3.98, p=0.023** | **$F_{(2,68)}$=4.02, p=0.022** | **$F_{(2,68)}$=3.93, p=0.024** |
| Offline | **$F_{(2,68)}$=4.33, p=0.017** | **$F_{(2,68)}$=4.38, p=0.016** | **$F_{(2,68)}$=4.17, p=0.020** |

*Statistical output assessing differences among the experimental groups in total, online and offline performance gains when separately controlling for the potential confounds of age, gender and education. The results presented in the main text remained significant when confounding variables were included as covariates in the statistical models. Thus, group differences in total, online and offline performance gains cannot be attributed to age, gender or education levels.*

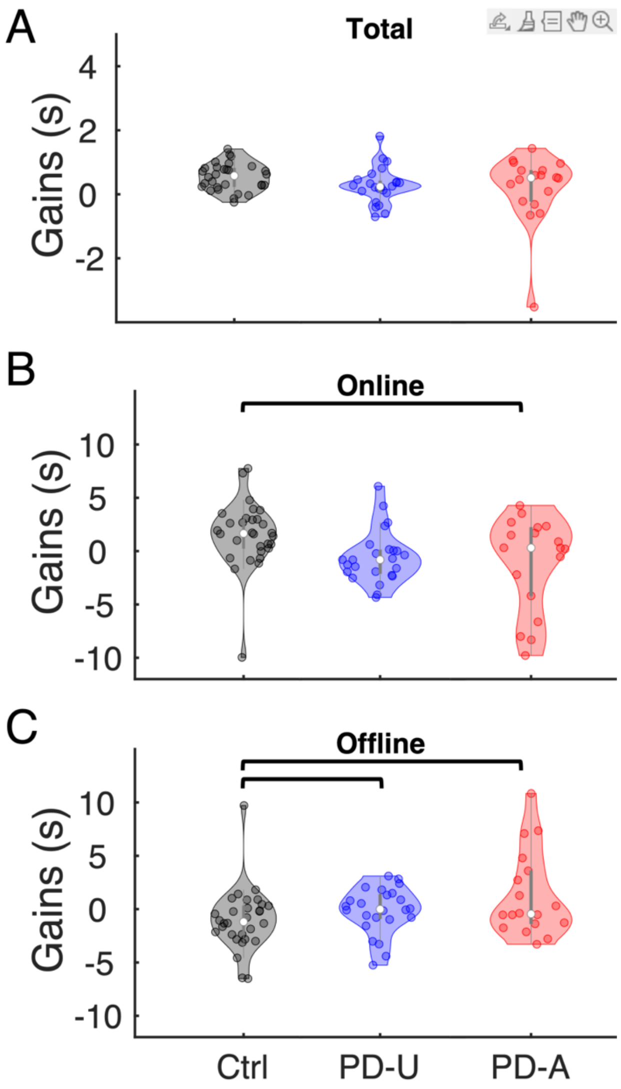


*Figure 2-1. Total (Panel A), micro-online (Online, B) and micro-offline (Offline, B) performance gains during the 24-hour retest session for the control (black; n=30), PD-U (blue; n=24) and PD-A (red; n=18) groups. Shaded regions represent kernel density estimates of the data, colored circles depict individual data, open circles represent group medians and vertical gray lines show interquartile ranges (Bechtold, 2016). There were no differences in total performance gains among the three groups ($F_{(2,69)}$=1.38, p=0.26, ges=0.038, $BF_{10}$=0.33), a result that was consistent with the data from the initial training session (see Figure 2 in the main text). A Group (3 levels) x Interval (2 levels: micro-online vs. -offline) ANOVA on performance gains revealed a significant interaction that again mirrored the results from initial training ($F_{(2,69)}$=3.68, p=0.030, ges=0.095, $BF_{10}$=42.94). Follow-up simple effects revealed that the three groups differed in the magnitude of both micro-online and -offline performance gains (online: $F_{(2,69)}$=3.71, p=0.029, ges=0.097, $BF_{10}$=1.97; offline: $F_{(2,69)}$=3.59, p=0.033, ges=0.094, $BF_{10}$=1.82). Pairwise comparisons revealed that the PD-A group exhibited significantly smaller and larger micro-online and micro-offline*

*performance gains, respectively, as compared to the controls (online: $t(46)=2.42$, $p=0.019$, Hedges $g=0.71$, $BF_{10}=2.93$; offline: $t(46)=2.32$, $p=0.025$, Hedges $g=0.68$, $BF_{10}=2.43$). The PD-U group also exhibited significantly larger micro-offline gains as compared to controls $t(52)=2.04$, $p=0.047$, Hedges $g=0.55$, $BF_{10}=1.49$), yet the difference in micro-online gains was considered as non-significant trend ($t(52)=1.91$, $p=0.061$, Hedges $g=0.51$, $BF_{10}=1.21$). Collectively, these results from the 24-hour retest session are analogous to those from initial training, with the exception that the control and PD-U groups did not exhibit a significant difference in the magnitude of micro-online performance gains in the 24-hour retest session. See Extended Data Table 2-3 below for details from within-group statistical contrasts.*

*Table 2-3. Within-group statistical results from the 24-hour retest session.*

| | Ctrl | PD-U | PD-A |
|---|---|---|---|
| Total | **$t_{(29)}$=6.98, p<0.001** | **$t_{(23)}$=4.16, p<0.001** | $t_{(17)}$=0.80 p=0.43 |
| Online | **$t_{(29)}$=2.70, p=0.011** | $t_{(23)}$=0.04, p=0.96 | $t_{(17)}$=1.05, p=0.31 |
| Offline | $t_{(29)}$=1.89, p=0.068 | $t_{(23)}$=1.02, p=0.32 | $t_{(17)}$=1.39, p=0.18 |

*Statistical output of one-sample t-tests assessing whether the magnitudes of total, micro-online (Online) and micro-offline (Offline) gains in performance from the 24-hour retest session differed from a test value of 0. Bold = significant at $p < 0.05$. Whereas the control and PD-U groups exhibited significant and positive total performance gains, suggesting continued improvements in the 24-hour retest session, the PD-A group did not. Performance improvements in the control group could largely be attributed to the positive online gains that significantly differed from the test value of 0.*

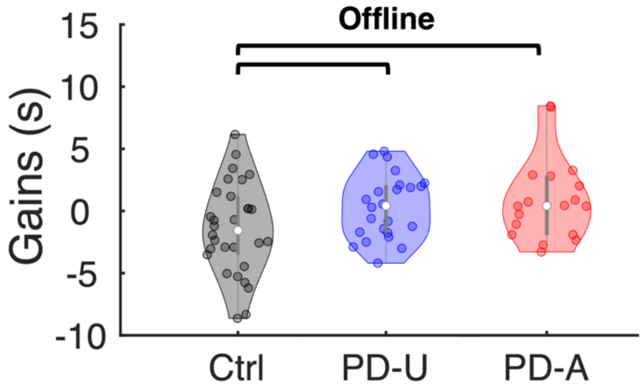


*Figure 2-2. Micro-offline performance gains – re-computed to minimize the effects of fatigue - for the control (black), PD-U (blue) and PD-A (red) groups. In contrast to data depicted in Figure 2C in the main text, micro-offline gains were the differences between the fastest – as opposed to the last - Sequence Duration in block n and the first Sequence Duration in block n+1. This computation thus minimizes the effects of progressively worsening performance within a block due to factors such as fatigue, inattention, etc. Positive values are indicative of performance improvements. Shaded regions represent kernel density estimates of the data, colored circles depict individual data, open circles represent group medians and vertical gray lines show interquartile ranges (Bechtold, 2016). Brackets: $p < 0.05$. The PD-U and the PD-A groups exhibited significantly larger micro-offline gains as compared to the controls (PD-U vs. controls: $t_{(52)}=2.59$, $p=0.013$, FDR-corrected $p=0.038$, Hedges $g=0.70$, $BF_{10}=4.01$; PD-A vs. controls: $t_{(46)}=2.27$, $p=0.028$, FDR-corrected $p=0.042$, Hedges $g=0.66$, $BF_{10}=2.22$; PD-U vs. PD-A: $t_{(40)}=0.09$, $p=0.93$, Hedges $g=0.03$, $BF_{10}=0.31$). This suggests that the larger micro-offline gains observed in individuals with PD cannot be explained by a disproportionately larger accrual of fatigue, inattention, etc. over online intervals that then dissipate during the micro-offline epochs.*

### Extended Data References

Bechtold B (2016) Violin Plots for Matlab. Github Project https://github.com/bastibe/Violinplot-Matlab, DOI: 105281/zenodo4559847.
